# HuR inhibition reduces post-ischemic cardiac remodeling by dampening acute inflammatory gene expression and the innate immune response

**DOI:** 10.1101/2023.01.17.524420

**Authors:** Samuel Slone, Sarah R. Anthony, Lisa C. Green, Michelle L. Nieman, Perwez Alam, Xiaoqing Wu, Sudeshna Roy, Jeffrey Aube, Liang Xu, John N. Lorenz, A. Phillip Owens, Onur Kanisicak, Michael Tranter

## Abstract

Myocardial ischemia/reperfusion (I/R) injury and the resulting cardiac remodeling is a common cause of heart failure. The RNA binding protein Human Antigen R (HuR) has been previously shown to reduce cardiac remodeling following both I/R and cardiac pressure overload, but the full extent of the HuR-dependent mechanisms within cells of the myocardium have yet to be elucidated. In this study, we applied a novel small molecule inhibitor of HuR to define the functional role of HuR in the acute response to I/R injury and gain a better understanding of the HuR-dependent mechanisms during post-ischemic myocardial remodeling. Our results show an early (two hours post-I/R) increase in HuR activity that is necessary for early inflammatory gene expression by cardiomyocytes in response to I/R. Surprisingly, despite the reductions in early inflammatory gene expression at two hours post-I/R, HuR inhibition has no effect on initial infarct size at 24-hours post-I/R. However, in agreement with previously published work, we do see a reduction in pathological remodeling and preserved cardiac function at two weeks post-I/R upon HuR inhibition. RNA-sequencing analysis of neonatal rat ventricular myocytes (NRVMs) at two hours post-LPS treatment to model damage associated molecular pattern (DAMP)-mediated activation of toll like receptors (TLRs) demonstrates a broad HuR-dependent regulation of pro-inflammatory chemokine and cytokine gene expression in cardiomyocytes. We show that conditioned media from NRVMs pre-treated with HuR inhibitor loses the ability to induce inflammatory gene expression in bone marrow derived macrophages (BMDMs) compared to NRVMs treated with LPS alone. Functionally, HuR inhibition in NRVMs also reduces their ability to induce endocrine migration of peripheral blood monocytes *in vitro* and reduces post-ischemic macrophage infiltration to the heart *in vivo.* In summary, these results suggest a HuR-dependent expression of pro-inflammatory gene expression by cardiomyocytes that leads to subsequent monocyte recruitment and macrophage activation in the post-ischemic myocardium.

## INTRODUCTION

Myocardial ischemia/reperfusion (I/R) injury results in cardiomyocyte cell death and an acute loss of contractile function. The cardiac remodeling that subsequently occurs in compensatory response to I/R is a common underlying cause of heart failure. Specifically, post-ischemic remodeling is known to be driven in part by an increase in the expression of cytokines and chemokines within the ischemic and ischemic border zones of the heart that elicit the innate inflammatory response.^1–3^ Within the first few hours to days, apoptotic and necrotic cell death results in the release of damage associated molecular patterns (DAMPs) that trigger pro-inflammatory signaling through stimulation of toll-like receptors (TLRs) on surviving cells.^1–3^ TLR receptor stimulation then leads to the expression and secretion of critical pro-inflammatory cytokines such as TNF-α, IL-6, and IL-1β.

The RNA binding protein Human Antigen R (HuR) is a widely expressed RNA binding protein that is an established mediator of inflammatory cytokine expression.^4^ Indeed, HuR expression has been shown to be increased at three days post-MI in a mouse model of non-reperfused ischemia.^5^ This same group also showed that HuR inhibition was able to reduce pathological remodeling and preserve cardiac function in this model, in part through abrogation of direct HuR-mediated expression of p53 and TGF-β in macrophages.^6^ However, many questions remain regarding the functional and mechanistic role of HuR in the acute response to I/R injury.

Thus, the primary goals of this work are to show the functional impact of HuR in mediating the acute expression of pro-inflammatory cytokines within the first few hours of cardiac reperfusion and subsequent effect on the initial myocardial infarct size at 24-hours post-I/R. To achieve this, we utilized a novel small molecule pharmacological inhibitor of HuR, KH-3, that has been shown to directly compete with HuR-RNA binding in a safe and specific manner *in vivo*^7,8^ Our results show that HuR inhibition elicits a robust decrease in acute inflammatory gene expression in cardiomyocytes within the first few hours following I/R that, surprisingly, has no effect on the initial infarct or myocyte cell death at 24-hours post-I/R. However, in agreement with aforementioned prior work, HuR inhibition does mitigate post-ischemic pathological remodeling and fibrosis and results in improved cardiac function at two weeks post-I/R. An unbiased RNA-sequencing approach using neonatal rat ventricular myocytes (NRVMs) treated with a TLR agonist with or without KH-3-mediated HuR inhibition demonstrates an overwhelming majority of HuR-dependent pro-inflammatory genes in response to TLR signaling in cardiomyocytes. Mechanistically, we then show that this HuR-dependent inflammatory gene expression in cardiomyocytes mediates the endocrine recruitment and activation of monocytes/macrophages. In sum, this work demonstrates that HuR inhibition provides cardioprotection from post-ischemic remodeling by reducing pro-inflammatory gene expression in cardiomyocytes and subsequent activation of the innate immune response.

## METHODS

### Ischemia/Reperfusion (I/R) Surgery

10–16-week-old male and female C57BI/6 mice were obtained from Jackson Labs (stock #000664). I/R surgery was performed as previously described.^9,10^ Briefly, mice were anesthetized under isoflurane, intubated, and ventilated. Following thoracotomy, the left anterior descending (LAD) coronary artery was visualized and a suture passed under the LAD distal to the left atrial appendage, immediately after the bifurcation of the major left coronary artery. Ischemia was induced by tightening the suture, and confirmed visually by blanching of the ventricle and by electrocardiogram displaying an elevated ST-segment. Just prior to loosening of the ligation, mice were randomly administered either VEH (5% EtOH, 5% Tween 80 and ddH20) or a small molecular inhibitor of HuR denoted as KH-3 (dosed at 60mg/kg up to 3x weekly). After 30 minutes, the suture was released to re-establish blood flow, and the incision closed in layers. Sham surgeries were performed following the same protocol but without tightening the suture of the left anterior descending coronary artery.

### Pharmacological Inhibition of HuR

HuR was pharmacologically inhibited using the small molecule KH-3, a second generation of HuR inhibitor compounds previously described by Wu et al.^11^ *In vivo* HuR inhibition was achieved via i.p. injection of KH3 3× weekly at a dose of 60 mg/kg as we have done previously^7^. *In vitro* HuR inhibition was achieved using KH-3 at a concentration of 20 μm.

### Echocardiography

All echocardiographic studies were performed as previously described.^7,12,13^ Briefly, mice were anesthetized under isoflurane and laid in a supine position on a heated platform with ECG recording. Body temperature was monitored and maintained at 36–38°C. Hair was removed from the chest using chemical hair remover or an electric razor prior to echocardiography. Using a Vevo 2100, parasternal images were obtained in short and long axes in two-dimensional mode and motion (M)-mode for quantification. These images were then analyzed using VevoStrain software (Vevo 2100, v1.1.1 B1455, Visualsonic, Toronto, Canada) for quantification of cardiac function.

### Infarct analysis

Mice were anesthetized with isoflurane 24□h after ischemia/reperfusion surgery and triphenyltetrazolium chloride (TTC) staining was performed as previously described.^14–16^ Briefly, hearts were excised and washed in a heparin/PBS mixture (0.5ml 1000U Heparin/10ml PBS) followed by cannulation of the aorta and perfusion with PBS. The LAD was occluded as for I/R surgery and hearts perfused with 150-200ul of 1% Evans blue. Hearts were then frozen at −20 °C for 60 minutes and then sectioned into six continuous slices 1□mm thick from apex to the occlusion site. Heart sections were incubated in 1% triphenyltetrazolium chloride (TTC) at 37 °C for 20 min followed by a fixation with 10% buffered formalin for 1 hour. Images were taken and analyzed with the use of NIH image software (Image J) of infarct zone (white), risk region (red), and non-risk region (blue).

### Neonatal rat ventricular myocyte isolation and culture

NRVMs were isolated using collagenase digestion and adhesion differential from fibroblasts as described.^17,18^ Briefly, Sprague Dawley neonatal rats (1-3 days old, Charles River) were decapitated, and the hearts were isolated. Following removal of the atria, the ventricles were cut into small pieces and digested first in .05% trypsin/EDTA (Corning) overnight. The next day trypsin was aspirated, and hearts were placed in collagenase II (Gibco) for 15 minutes in a 37°C water bath followed by 15 minutes of incubation at 37°C, 5% CO2. Cells were manually dissociated by trituration and the supernatant passed through a 40um strainer. Cells were then spun at 100 × g followed by a 40-minute pre-plating process on non-treated plates to allow for fibroblasts adherence. The nonadherent NRVMs were then transferred to gelatin (0.1%) coated culture-treated dishes in MEM alpha media (Gibco) with 10% FBS and 1% Penicillin-Streptomycin. NRVMs were treated with lipopolysaccharide (LPS) at a concentration of 1ug/ml and inhibition of HuR was achieved via siRNA (see below) or via small molecular inhibitor KH3 at a concentration of 20um.

### Bone marrow-derived macrophage (BMDM) isolation and culture

Wild-type C57BI/6 mice were sacrificed using CO_2_ and subsequent cervical dislocation followed by removal of the femurs and tibias. BMDM were then isolated as described^19,20^ with slight modifications. Briefly, once femurs and tibias were cut out and cleaned with ethanol, the end of the bone with visibly less marrow was cut off and the bones were placed (cut end down) into a 0.5mL tube with a hole in the bottom, which was then placed in a larger (1.5ml) tube. Quick pulses for 4-5 seconds were performed until all or most of the red bone marrow was collected in the outer tube. The collected red bone marrow was distributed into DMEM media (20% FBS and 1% P/S) containing 50ng/ml of Macrophage colony-stimulating factor (M-CSF) in 100cm petri dishes and kept in 37°C, 5% CO2 incubator. Post 7 days of plating, non-adherent cells were discarded, and BMDMs were re-plated in RPMI media containing 10% FBS and 1% P/S and plated at a density 1.2×10^6^ per 1.5ml for a 6 well plate.

### siRNA-mediated gene silencing

To achieve siRNA-mediated knockdown of HuR expression, NRVMs were seeded at □75% confluency and transfected with HuR or nontargeting control siRNA (80 nM) (Santa Cruz Biotechnology) 24 hours after plating using Lipofectamine 3000 (ThermoFisher Scientific) as per manufacturer’s instructions. Cells were grown for 48 hours post-transfection prior to further experimental treatment.

### RNA isolation and qPCR

RNA was isolated using the Macherey-Nagel NucleoSpin RNA kit. RNA quantity and quality was assessed by optical density at 260 nm and optical density ratios of 260/280 nm and 260/230 nm ratios, respectively. cDNA was synthesized using iScript Reverse Transcription Supermix for RT-qPCR (BioRad). Samples were run on CFX96 Touch Deep Well Real-Time PCR Detection System (BioRad) using iTAQ Universal SYBR Green qPCR Master Mix (BioRad) to assess levels of GAPDH, 18s, IL-6 (interleukin-6), TNF-alpha (Tumor Necrosis Factor alpha), IL-1 alpha (Interleukin 1 alpha), IL-1 beta (Interleukin 1 beta). Results were analyzed using the ΔΔCt method.^21^ Primers are as listed:

*18s:* F: 5’–AGTCCCTGCCCTTTGTACACA-3’, R: 5’–CCGAGGGCCTCACTAAACC-3’
*IL-6:* F: 5’–CCGGAGAGGAGACTTCACAG-3’, R: 5’–TGCCATTGCACAACTCTTTTCTC-3’
*TNF-alpha:* F: 5’–CCCATATACCTGGGAGGAGTCTTC-3’, R: 5’CATTCCCTTCACAGAGCAATGAC-3’,
*IL-beta:* F: 5’–AACCTGCTGGTGTGTGACGTTC-3’, R: 5’–CAGCACGAGGCTTTTTTGTTGT-3’.

### Protein Isolation and Western blotting

Total protein was isolated from *in vitro* cell cultures or heart tissue using RIPA Buffer with protease inhibitor cocktail. In addition, nuclear and cytoplasmic fractions were collected from in vivo heart tissue as previously described^22^. 15 ug of total protein, or a ratio of 10ug (nuclear)/30ug (cytoplasmic) extract, per lane was separated on a 10% polyacrylamide gel and transferred to a nitrocellulose membrane. Blocking was performed for 1 hour at room temperature using 5% bovine serum albumin in 0.1% Tween 20, tris-buffered saline (T-TBS) solution. Primary antibodies for HuR (1:2000), Periostin (1:1000), Histone H3 (1:1000), CD68 (1:500), GAPDH (1:2000) were incubated overnight at 4 °C in 5% BSA, and secondary antibodies (1:5000) were incubated for 1 hr. at room temperature in 5% BSA. Protein loading was normalized using Stain Free total protein, Histone H3 (nuclear) or GAPDH (cytoplasmic), as appropriate.

### Histological Analysis

Mouse hearts were isolated and embedded in Tissue-Tek OCT compound then rapidly frozen in an acetone and dry ice slurry. Frozen hearts were sectioned at 7 μm thickness and were subsequently post-fixed and processed for histological analysis. picrosirius red, sections were stained with Picrosirius Solution B (Polysciences, 24901B) for one hour, washed in 0.1M HCl, and then dehydrated and cleared through ethanol and xylene washes, according to the manufacturer’s protocol. Immunostaining of macrophages was performed using a CD68 antibody as previously described.^23^ Briefly, sections were incubated in blocking solution (5% BSA, Sigma Aldrich) for 30 minutes, followed by primary CD68 antibody for one hour and AlexaFluor secondary antibody (Thermo Fisher Scientific A11008) for 30 minutes. TUNEL staining was performed using a commercial kit (In Situ Cell Death Detection Kit, TMR (Sigma, 12156792910)) according to the manufacturer’s protocol. Briefly, frozen sections were post-fixed with paraformaldehyde followed by permeabilization (0.1% triton X-100), then incubated with TUNEL reaction mixture at 37 °C and nuclei were counterstained with DAPI. All images were captured on a BioTek Cytation 5 imager and analyzed in Image J. For

### RNA Sequencing and Gene Ontology Analysis

RNA was isolated from NRVMs as described above and 300 ng total RNA was sequenced using 100 base pair paired end reads at a total depth of approximately 40 million reads per sample. Genomic mapping of sequence reads and differential expression analysis, using a statistical threshold of a false discovery rate (FDR)-corrected P-value ≤ 0.01 and fold-change ≥ 2, was performed using the Qiagen CLC Genomics Workbench (version 21.0.3) as previously described by our lab.^13,24,25^ Principal component analysis (PCA), heat map, and Venn diagram were also all done using CLC Genomics Workbench. All RNA-seq raw data files and total gene count files have been deposited to the NCBI Gene Expression Omnibus (GEO) (https://www.ncbi.nlm.nih.gov/geo) (GSE Accession Number Pending).

Subsequently gene ontology (GO) analysis of differentially expressed genes was done using the NIH DAVID Bioinformatics Functional Annotation Tool (https://david.ncifcrf.gov) using a statistical threshold of FDR P-value ≤ 0.01 and ≥ 5 total genes per GO term. ^26,27^ GO enrichment plots were created using RStudio (version 2021.09.2+382) on macOS.

### PBMC Isolation and Migration Assay

Peripheral blood mononuclear cells (PBMCs) were isolated from the blood of adult rats as previously described.^28^ Briefly, PBMCs were obtained using the Ficoll-Paque centrifugation (GE Healthcare, 17–5446–02), according to the manufacturer’s instructions. Following isolation, PBMCs were plated into the inserts of the migration chamber at a concentration of 1×10^6 per well. The protocol for the migration assay is described by the manufacturer’s protocol (CellBioLabs).

### Statistical Analysis

Statistical analysis was performed using GraphPad PRISM software. All graphed results are presented as the mean ± standard deviation. Results were analyzed with unpaired Student’s t-tests, one-way ANOVA, or two-way ANOVA as appropriate, and statistical significance between groups was considered at P ≤ 0.05.

### Study Approval

All animal procedures were performed with the approval of the Institutional Animal Care and Use Committee of the University of Cincinnati and in accordance with the NIH Guide for the Care and Use of Laboratory Animals.

## RESULTS

### HuR is activated 2 hours post-I/R and total protein expression is increased 7 days post-I/R

Krishnamurthy *et al* previously showed that total HuR expression is increased at three days post-I/R and shRNA-mediated knockdown of HuR reduced inflammatory signaling at three days post-MI. ^5,6^ To determine if HuR is acutely activated in the infarcted myocardium, wild-type C57BI/6 mice were randomized to either 30 minutes of LAD occlusion or sham surgery followed by two hours reperfusion. Western blot analysis shows a significant increase in HuR nuclear-to-cytoplasmic translocation in the ischemic zone of I/R hearts compared to sham at two hours post-reperfusion (Fig.1A-B). In addition, total protein collected from the LV ischemic zone shows increased total HuR expression in I/R hearts compared to sham mice at seven days post-I/R (Fig. 1C-D). These results show that HuR is activated as soon as two hours post-I/R and total expression level remains increased for at least seven days.

**Figure 1.**
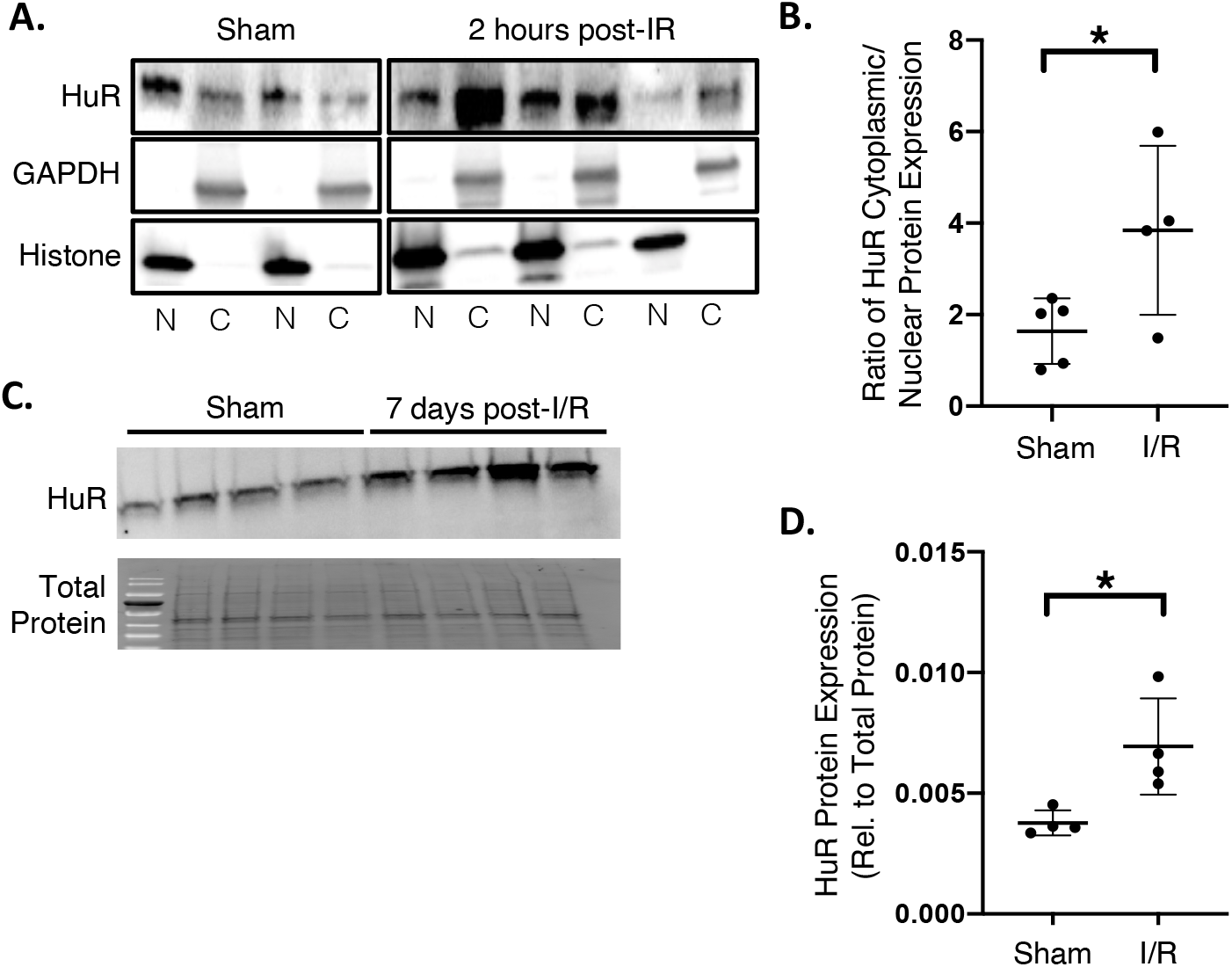
HuR activation and expression is increased following I/R injury. HuR cytoplasmic translocation, indicative of activation, is increased at two hours post-I/R (N and C represent nuclear and cytosolic fractions, respectively) (A). Cytoplasmic translocation of HuR is quantified in panel B. Total HuR protein expression is increased in hearts at seven days post-I/R (C, D). \**P* ≤ 0.05.

### HuR inhibition blunts cardiac cytokine expression, but does not impact acute infarct size

To determine the functional impact of HuR signaling in the acute response to I/R injury, mice were treated with a small molecule inhibitor of HuR (KH-3; HuRi), or vehicle control, at the time of reperfusion. Results show that HuR inhibition (HuRi) blunts the I/R-mediated increase in IL-6 and TNF-α gene expression in the infarct zone at two hours post-reperfusion (Fig. 2A-B). However, HuR inhibition had no effect on the initial infarct at 24 hours post-I/R at determined by TTC staining (Fig. 3A-B). In addition, TUNEL staining performed on cardiac sections spanning the infarcted regions of the showed no significant differences of the percentage of TUNEL positive cells between mice treated with vehicle and HuRi (Fig. 3C-D).

**Figure 2.**
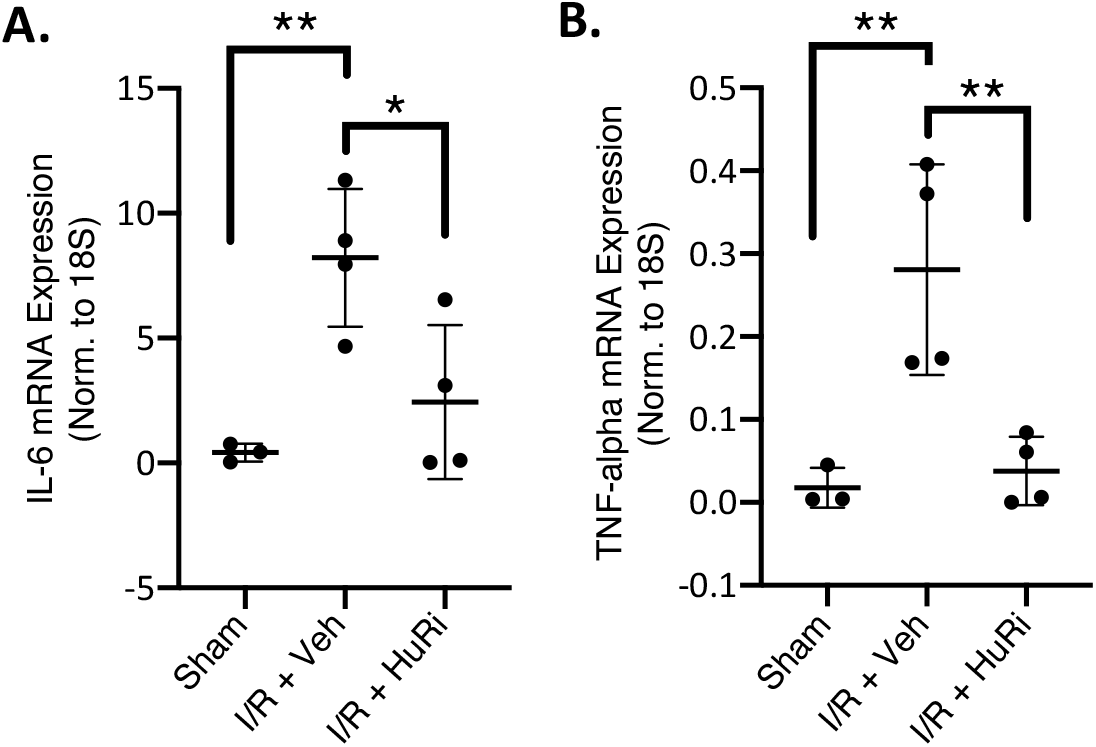
HuR inhibition reduces acute inflammatory gene expression post-I/R. Treatment with the HuR inhibitor KH-3 (50 mg/kg) at the time of reperfusion reduces immediate early expression of pro-inflammatory gene expression of IL-6 (A) and TNF-⍰ (B) at 2 hours postreperfusion. \**P* ≤ 0.05. \*\**P* ≤ 0.01.

**Figure 3.**
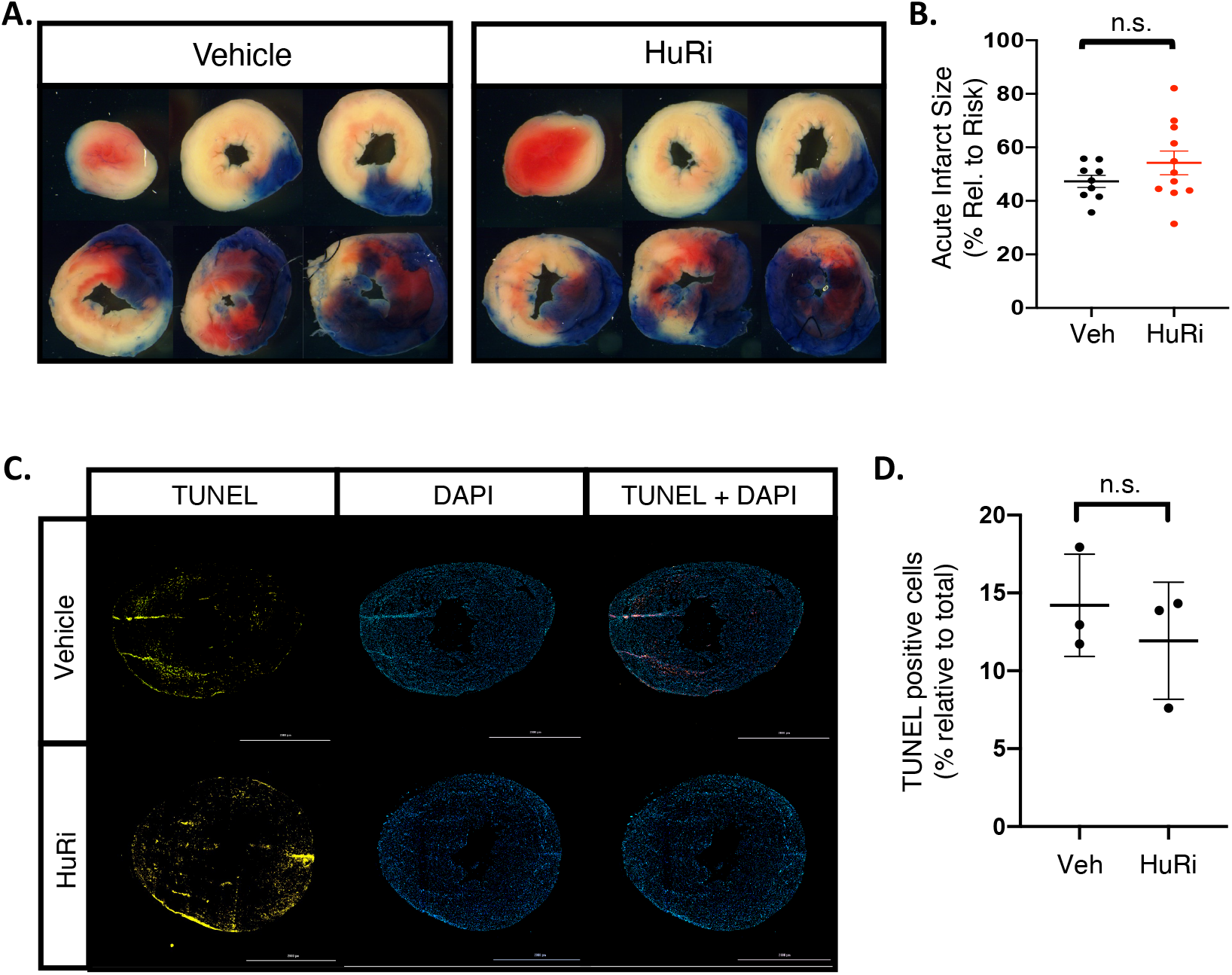
HuR does not mediate acute infarct size or I/R-mediated cell death. HuR inhibition has no impact on initial infarct size at 24 hours post-I/R as measured by TTC vital staining (A; quantified in B) or cell death as measured by TUNEL staining (C; quantified in D).

### HuR inhibition decreases post-ischemic ventricular remodeling and cardiac fibrosis

Cardiac structure and function were assessed via echocardiography at two weeks post-I/R to determine the functional effect of HuR inhibition. HuR inhibition blunts the post-ischemic increase in left ventricular (LV)-end diastolic and LV-end systolic volume at two weeks following I/R (Fig. 4A-B), while preserving cardiac ejection fraction (Fig. 4C) compared to vehicle treated mice. Additionally, mice that received HuR inhibitor displayed less total fibrosis at two weeks post-I/R as assessed by picrosirius red staining (Fig. 4D-E). Consistent with a reduction in total fibrosis, protein expression of periostin, an indicator of activated myofibroblasts, was also significantly decreased at two weeks post-I/R upon HuR inhibition (Fig. 4F-G).

**Figure 4.**
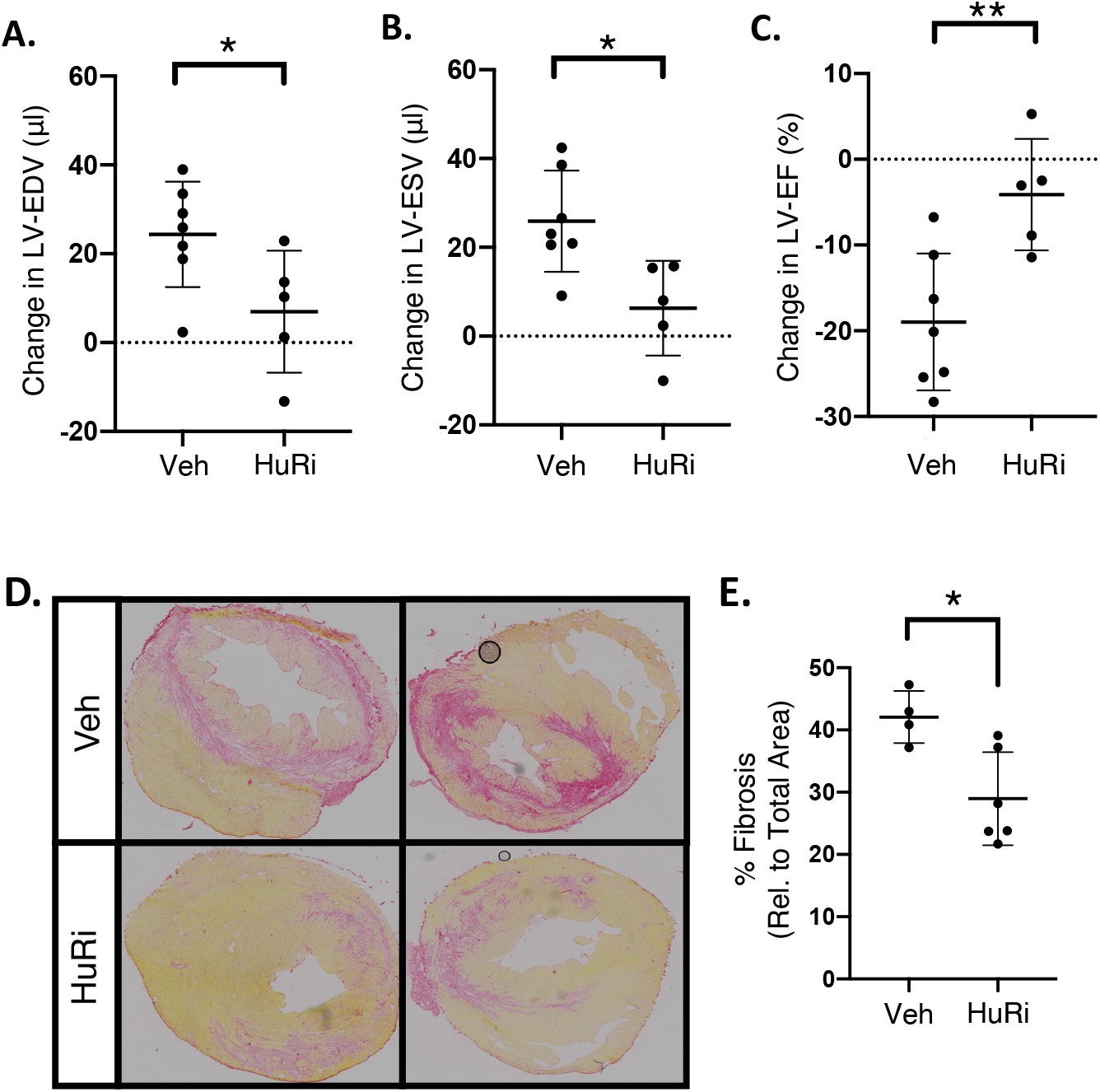
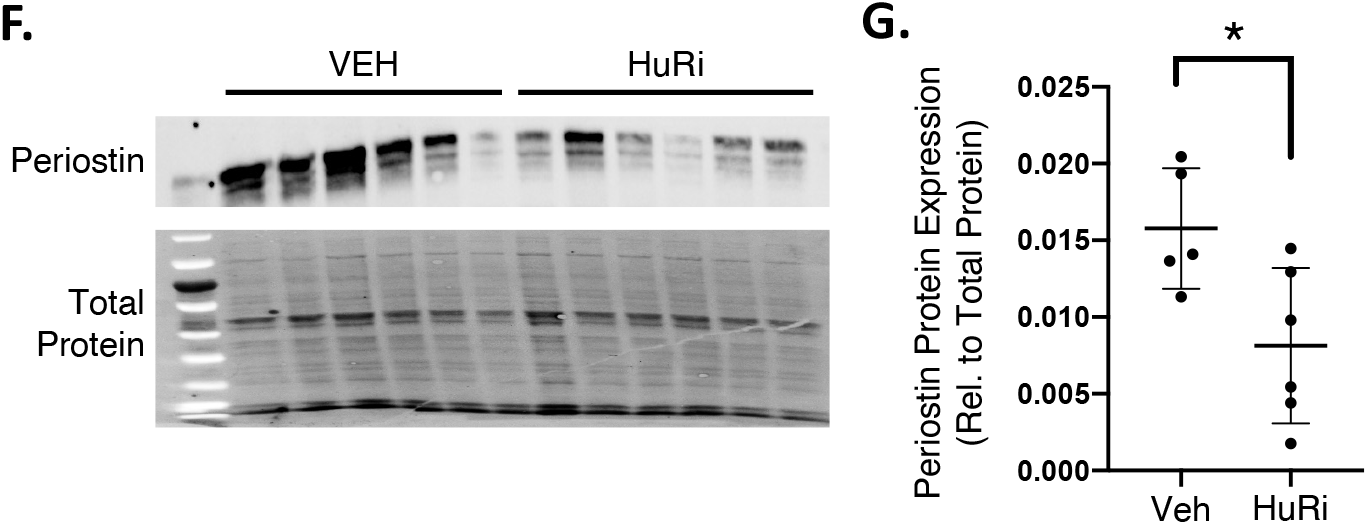
HuR inhibition reduces post-ischemic cardiac remodeling and preserves cardiac function. HuR inhibition significantly reduced post-ischemic cardiac remodeling as assessed by LV end diastolic volume (A) and end systolic volume (B) and preserved LV function (ejection fraction; C) at two weeks post-I/R. Treatment with HuRi also reduced total cardiac fibrosis as determined by picrosirius red staining (D; quantified in E) and myofibroblast activity as assessed by total expression of the myofibroblast marker periostin (F; quantified in G). \**P* ≤ 0.05. \*\**P* ≤ 0.01.

### HuR mediates the expression of inflammation and immune response related genes in myocytes following LPS stimulation

To begin to delineate the mechanisms of HuR-dependent effects in the post-ischemic myocardium, we utilized a model of cultured neonatal rat ventricular myocytes (NRVMs) treated with LPS to mimic toll-like receptor (TLR) receptor signaling via damage associated molecular patterns (DAMPs) that often occurs during *in vivo* I/R injury.^1–3^ NRVMs were treated with LPS with or without HuR inhibition and RNA-sequencing was performed at two hours post-treatment to mirror the *in vivo* timepoint at which HuR inhibition was shown to suppress I/R-induced IL-6 and TNF-α expression.

Principal component plot analysis (PCA; Fig. 5 A) and gene expression heat map (Fig. 5 B) were used to visualize broad level changes in the transcriptome and demonstrate clear changes in the transcriptome from NRVMs treated with LPS, LPS plus HuR inhibitor, or vehicle control for 2 hours. We next applied differential expression analysis using a statistical cutoff of an FDR P-value ≤ 0.01 and fold-change ≥ 2, and found 1911 total differentially expressed genes following LPS treatment (compared to vehicle), with 1113 of these LPS-dependent expression changes also dependent on HuR.

**Figure 5.**
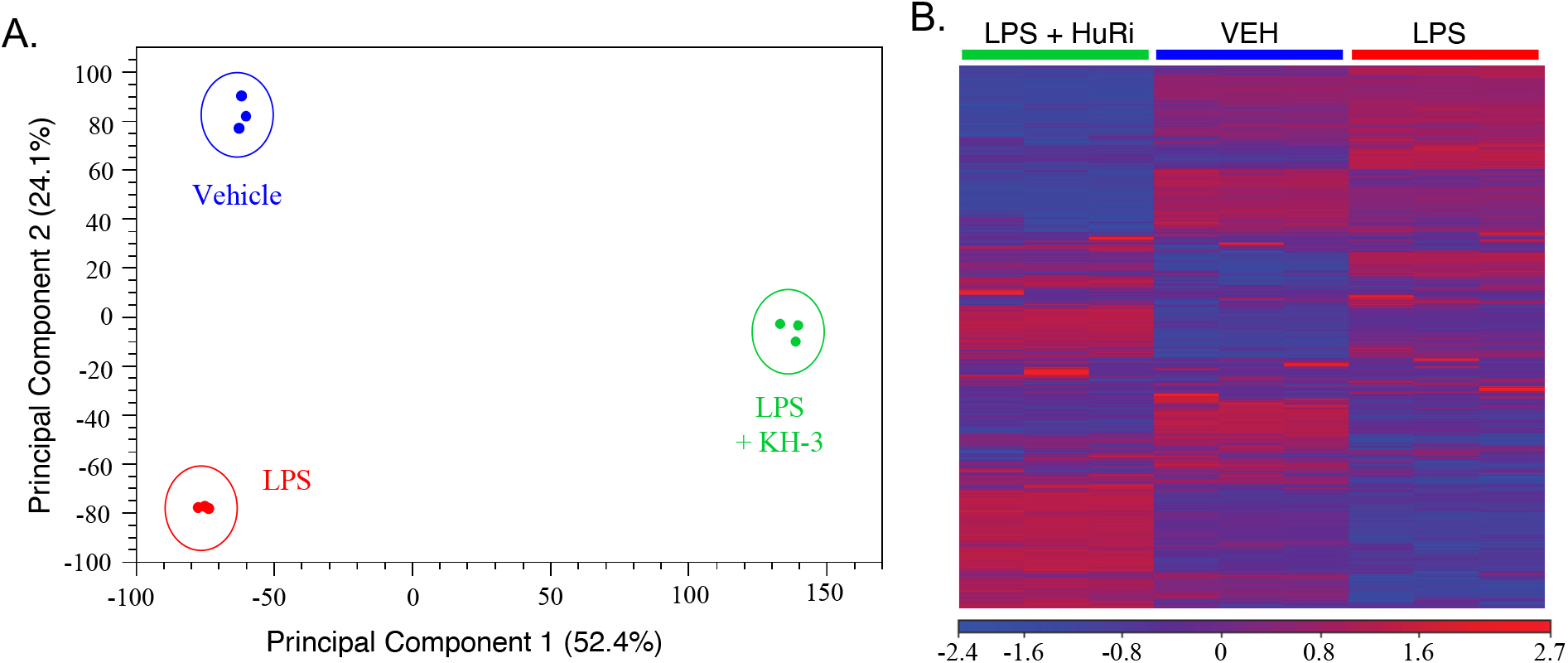
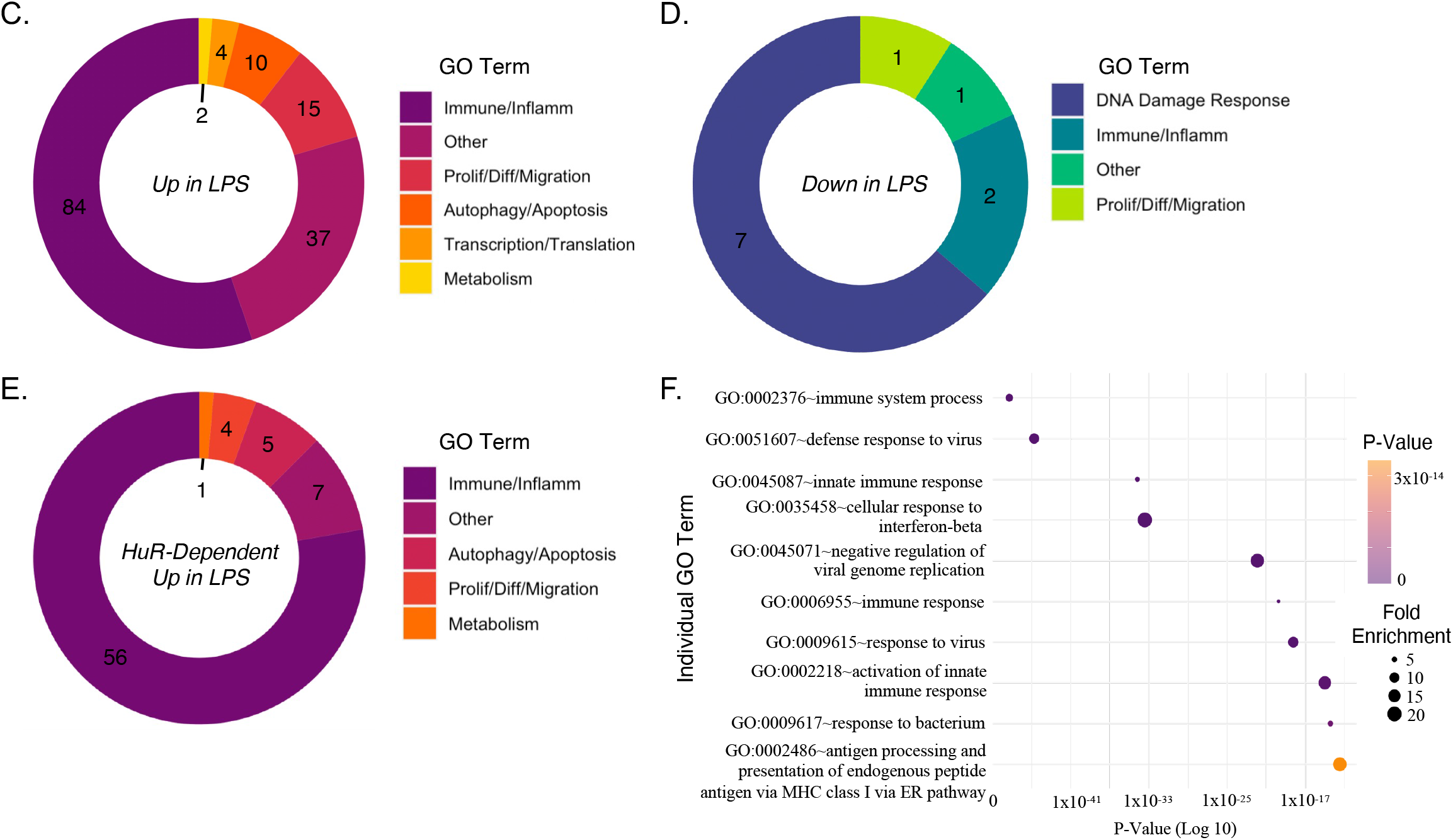
HuR mediates inflammatory gene expression following TLR stimulation in cardiomyocytes. Principle component analysis (PCA) (A) and heat map analysis (B) show a distinct and consistent differential gene expression profile in NRVMs treated with LPS alone or LPS plus HuRi (KH-3). Donut plot analysis of all LPS-dependent significant upregulated (C) and downregulated (D) gene expression changes grouped by gene ontology (GO) term. GO grouping of all HuR-dependent gene expression changes among those upregulated by LPS shows that HuR overwhelmingly regulates immune and inflammation related GO term-associated genes (E). The top ten HuR-dependent GO terms from panel E are graphed out by P-value and fold enrichment (F).

Gene ontology (GO) enrichment analysis was used to provide functional insight into these changes in gene expression and showed a significant enrichment in inflammation and innate immune response-related GO terms following LPS treatment (Fig. 5C). Unsurprisingly, the top ten enriched LPS-dependent GO terms among upregulated genes in myocytes, as ranked by P-value, are all related to inflammation or the innate immune response (Fig. S1). Conversely, only 11 GO terms were significantly enriched for among the LPS down-regulated genes, and these were predominately associated with the DNA damage response and DNA repair (Fig. 5D and S2).

Importantly, of the 72 significantly enriched HuR-dependent GO terms within genes whose expression was significantly increased following LPS stimulation, 56 (78%) of these were related to inflammation or the innate immune response, indicating these pathways to be primarily HuR-dependent downstream of TLR signaling in myocytes (Fig. 5E-F). On the other hand, among genes significantly downregulated following LPS stimulation, very few (only 3) of these significantly down-regulated GO enrichments were found to be HuR-dependent (Fig. S3).

These results, showing that HuR primarily mediates increased gene expression following LPS treatment, are not surprising given the primary role of HuR in stabilizing, and thus promoting, increased expression of mRNA targets. It is also worth noting that the majority of the significant LPS-dependent gene expression changes (1040/1911; 54%) were LPS-induced increases in expression (Table S1), and 617 of these (59%) were found to also be HuR-dependent (Table S2). Taken together, these results suggest that HuR plays a significant role in the LPS/TLR-dependent increases in inflammatory and innate immune system related gene expression in myocytes.

### Cardiomyocytes induce inflammatory gene expression in macrophages via a HuR-dependent endocrine mechanism

Since HuR-dependent gene expression in myocytes appears to be independent of myocyte cell death or acute infarct size, we next sought to determine if these HuR-dependent gene expression changes in myocytes might have an impact on post-ischemic remodeling through endocrine effects to other cell types. Recruitment and activation of the innate immune response is known to play a substantial role in post-ischemic cardiac remodeling.^1–3^ Indeed, we observe a decrease in macrophage infiltration to the myocardium at seven days post-I/R upon HuR inhibition (Fig. 6A-B).

**Figure 6.**
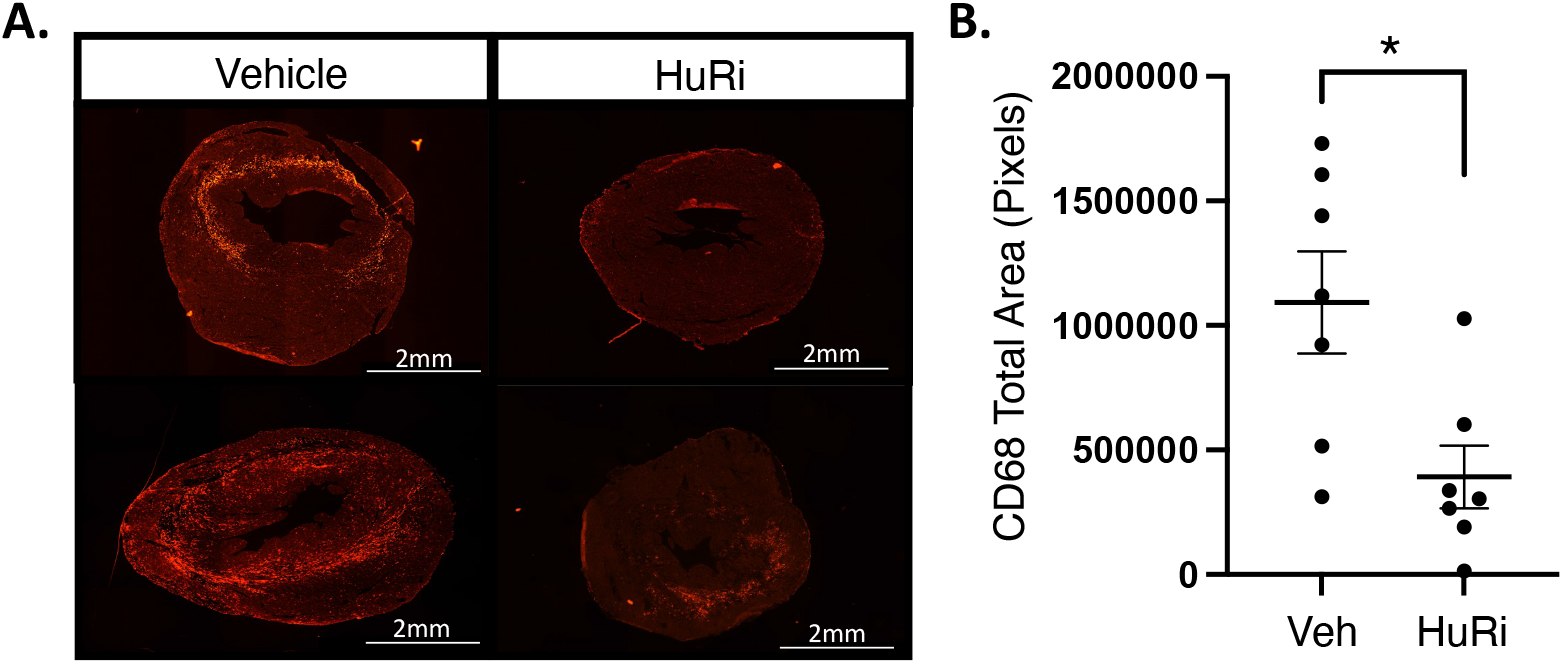
HuR inhibition reduces macrophage infiltration to the post-ischemic myocardium. Immunofluorescent staining of CD68^+^ cells shows a reduction in macrophage infiltration to the heart at seven days post-I/R upon HuR inhibition (A; total area of CD68^+^ stain is quantified in B). \**P* ≤ 0.05.

Accordingly, we utilized a conditioned media transfer model from NRVMs to bone marrow derived macrophages (BMDMs) to determine if acute HuR-dependent signaling in myocytes mediates macrophage activity (Fig. 7A). HuR inhibition was confirmed by qPCR to decrease the LPS-induced expression of IL-1β and TNF-α in NRVMs (Fig. 7B-C). Following an acute (one hour) treatment with LPS, NRVMs were washed with PBS, and fresh media was added for 12 hours prior to transfer to naïve BMDMs. Results show TNF-α gene expression in BMDMs is significantly blunted when exposed to conditioned media from LPS + HuRi treated NRVMs compared to media from NRVMS treated with LPS alone (Fig. 7D). A similar decrease in BMDM TNF-α gene expression is observed when treated with conditioned media from NRVMs treated with LPS + HuR siRNA compared to LPS alone, validating the specificity of HuR pharmacological inhibition in this model (Fig. 7E-G).

**Figure 7.**
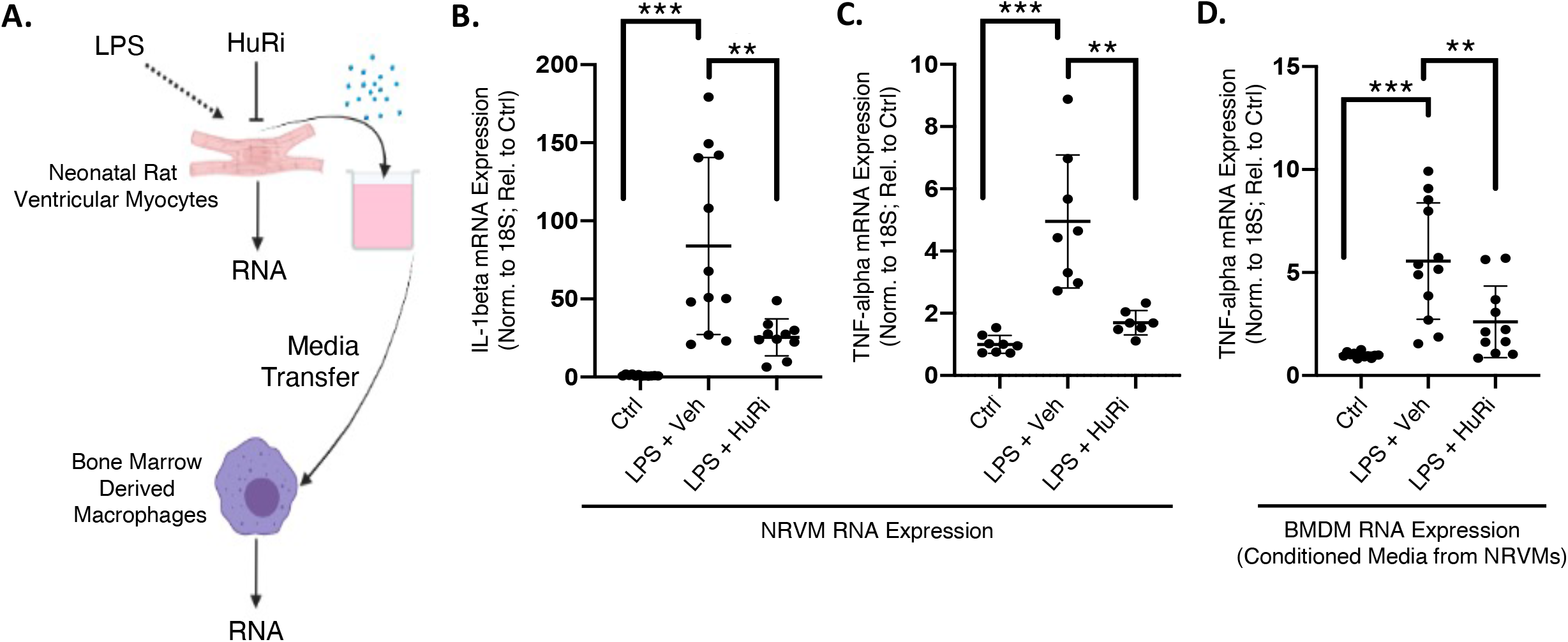
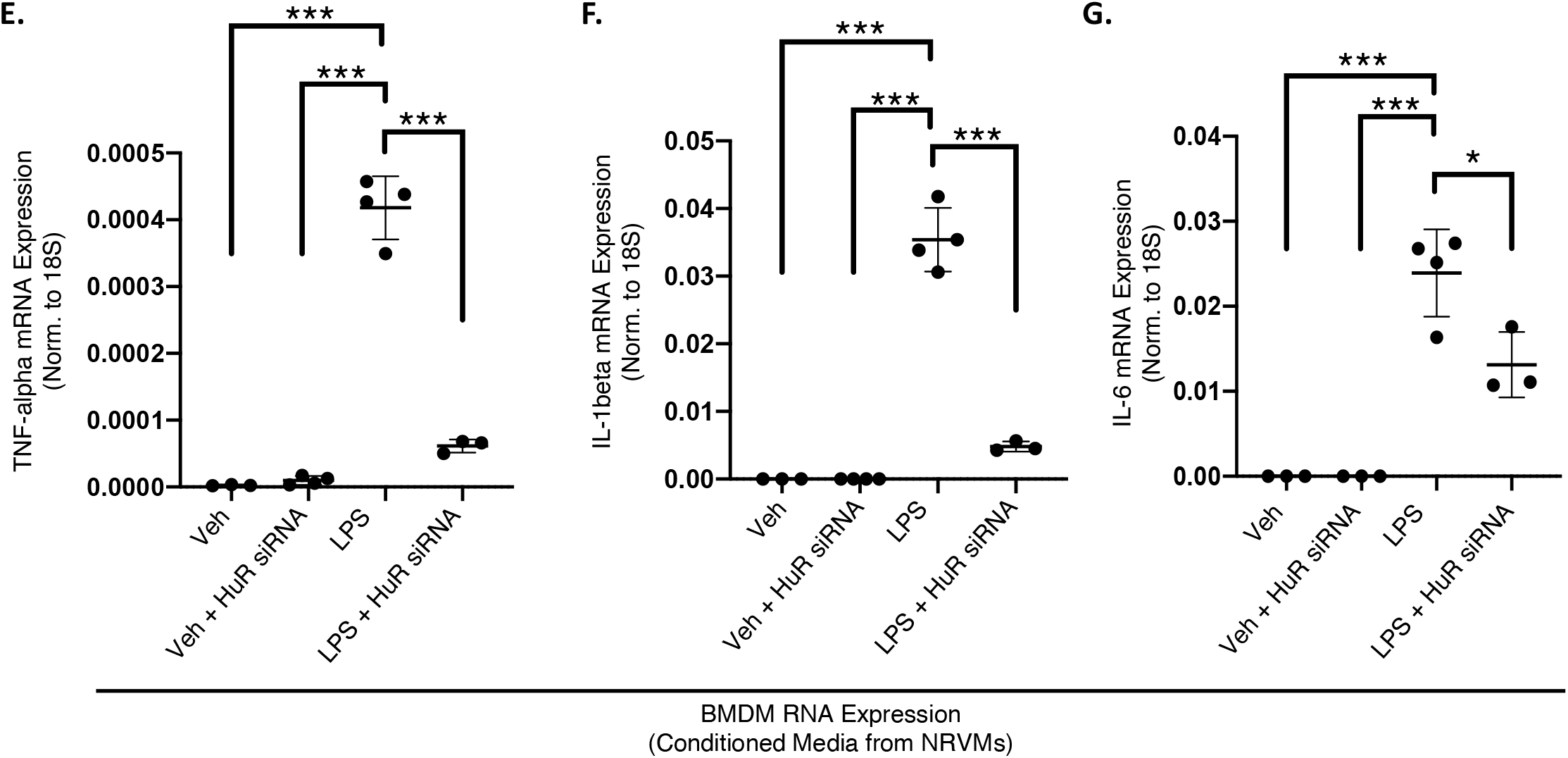
TLR signaling in myocytes induces inflammatory gene expression in macrophages in a HuR-dependent endocrine manner. Schematic representation of NRVM treatment and conditioned media transfer to BMDMs (A). HuR inhibition reduces mRNA expression of IL-1β (B) and TNF-α (C) in NRVMs. Conditioned media from LPS treated NRVMs induces expression of TNF-α in BMDMs that is blunted when NRVMs are treated with HuR inhibitor (D). siRNA-mediated knockdown of HuR in NRVMs inhibits the LPS-mediated increase in TNF-α (E), IL-1β (F), and IL-6 (G) in BMDMs treated with conditioned media from NRVMs. \**P* ≤ 0.05. \*\**P* ≤ 0.01. \*\*\**P* ≤ 0.001.

Importantly, we next utilized a peripheral blood mononuclear cell (PBMC) migration assay to determine whether the HuR-dependent endocrine signaling from myocytes is sufficient to induce functional changes in the migratory nature of these cells to the myocardium following ischemic injury. NRVMs were again treated with LPS alone or LPS plus HuRi and conditioned media was transferred to the top half of a migration chamber containing isolated PBMCs. Results demonstrate increased PBMC migration when exposed to conditioned media from NRVMs treated with LPS alone, but this migratory effect is lost upon HuR inhibition in the NRVMs (Fig. 8B).

**Figure 8.**
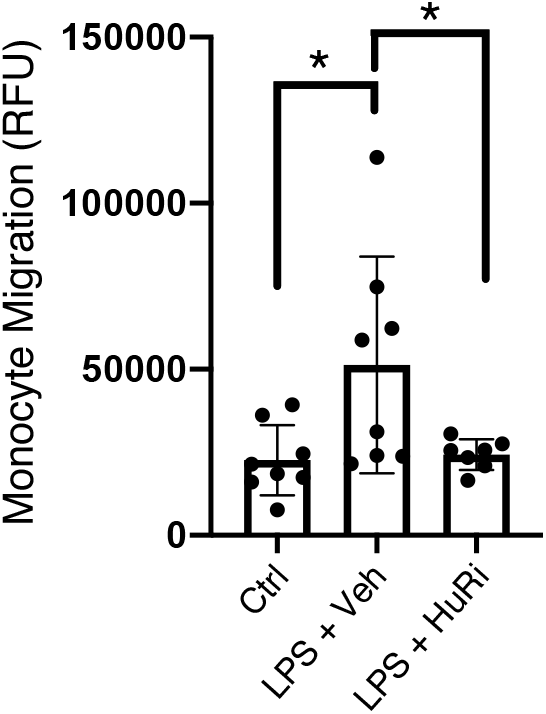
TLR signaling in myocytes induces monocyte migration in a HuR-dependent endocrine manner. Conditioned media transfer from NRVMs treated with LPS induces monocyte (PBMC) migration that is lost when NRVMs are pre-treated with HuR inhibitor. \**P* ≤ 0.05.

## DISCUSSION

The results presented herein demonstrate that pharmacological inhibition of HuR reduces pathological remodeling and preserves cardiac function in a mouse model of I/R injury. Furthermore, HuR inhibitor was given at the onset of cardiac reperfusion (following a 30-minute ischemia) mimicking a clinically relevant timepoint that may be achievable in a human study should an approved inhibitor of HuR become available. Indeed, HuR has been well-studied as an oncogene in multiple human cancers and the clinical development of HuR inhibitors are being actively sought as cancer therapeutics.^8,29–33^ Not as much is known regarding the functional role of HuR in cardiovascular diseases, but our lab and others have shown that HuR expression is increased in failing human hearts.^7,34^ Our previous work applied the same pharmacological inhibitor used in this study to show that HuR inhibition reduces pathological cardiac remodeling in a mouse model of chronic pressure overload.^7^ Chronic cardiac pressure overload (e.g. hypertension or aortic stenosis) and ischemic injury are two of the most common underlying causes of heart failure, but the molecular mechanisms by which each progresses are distinctly at the cellular level.

Previous work showed that administration of the anti-inflammatory cytokine IL-10 mitigates cardiac remodeling and dysfunction following chronic cardiac ischemia in a permanent ligation myocardial infarction (MI) model, in part through suppression of HuR expression.^5^ This same group subsequently showed that direct knockdown of HuR via shRNA yielded a similar reduction in remodeling and preserved function.^6^ However, while they did show elevation of total HuR protein levels as early as three days post-MI^5^, the acute effects of HuR-dependent signaling in the early response to ischemia (or reperfusion) were not addressed. Indeed, our results show that total HuR expression is elevated at seven days post-I/R, but we also show that nuclear-to-cytoplasmic translocation of HuR, indicative of HuR activation, is increased as early as just two hours after reperfusion. This is an important consideration not only for how the initial ischemic infarct size may affect subsequent remodeling (e.g. a smaller initial cardiac infarct size may mask a differential chronic remodeling effect), but also for mechanistic understanding of how HuR mediates the observed changes in cardiac structure and function.

We initially hypothesized that HuR inhibition would result in decreased inflammatory gene expression in the acute phase (0-24 hours post-I/R) and a subsequently smaller initial cardiac infarct size at 24-hours post-I/R. Thus, while our results showing reduced pathological remodeling and preserved function at two weeks post-I/R corroborate previously published work, our results showing no effect of HuR inhibition on acute infarct size were surprising. Despite this, HuR inhibition did blunt early (two hour post-I/R) inflammatory gene expression *in vivo,* a result that we were able to replicate using a direct LPS-mediated stimulation of TLRs in NRVMs *in vitro.* Application of RNA sequencing in NRVMs treated with LPS with and without HuR inhibition also showed that the overwhelming majority of HuR-dependent genes in these cells following LPS stimulation associate with inflammation-related gene ontology categories. Many of these HuR-dependent inflammatory gene products are secreted cytokines and chemokines that would exert likely paracrine or endocrine effects on other cells.

Our results show that HuR inhibition significantly reduces the number of CD68^+^ macrophages in the myocardium at seven days post-I/R. Our *in vitro* data using conditioned media from NRVMs treated with HuRi supports a HuR-dependent, cardiomyocyte-driven endocrine recruitment and/or activation of monocytes/macrophages as a likely mechanism. Deeper mechanistic studies, including inhibition of specific cytokine signaling within the recipient macrophages, will be needed to conclusively identify the responsible secreted factor, but our results clearly show that one or more HuR-dependent factors are secreted from NRVMs following TLR stimulation that are sufficient to induce inflammatory signaling in macrophages. Importantly, not only are these HuR-dependent secreted factors capable of activating macrophages, which may be most relevant to the existing pool of cardiac macrophages, but they are capable of inducing migration in naïve PBMCs. Functionally, this HuR-dependent early inflammatory cytokine and chemokine gene expression in myocytes from the ischemic myocardium may then contribute to the systemic innate immune response leading to a recruitment of monocytes to the injured myocardium and induction of pro-inflammatory gene networks in the cardiac macrophage population.

Interestingly, our previous work identified HuR-dependent regulation of TGFβ expression in myocytes as a likely paracrine cell-to-cell crosstalk mechanism of cardiac fibroblast activation during pressure overload-induced remodeling.^7^ We have since gone on to show that HuR is also highly expressed in fibroblasts and is a critical mediator of cardiac myofibroblast activation in response to TGFβ^35^, while other recent work has suggested HuR activity in macrophages to mediate fibrosis and inflammation in diabetic cardiomyopathy.^34^ Both of these works raise the independent possibilities that some of our observed protection here may be due to HuR inhibition in cardiac fibroblasts or macrophages in addition to cardiomyocytes.

Together, the existing literature is making a strong case for HuR as a central player in cardiac remodeling with diverse roles in multiple types. However, while a detrimental effect of HuR inhibition on the myocardium has yet to be identified – we previously showed no deleterious cardiac phenotype in myocyte-specific deletion or pharmacological inhibition of HuR for seven weeks in sham mice^7^ – additional work is needed to fully elucidate the function of HuR in other cardiac cell types. As such, HuR inhibition appears to be a promising therapeutic approach, dampening the pro-inflammatory and pro-fibrotic environment of the myocardium to reduce pathological remodeling in multiple etiologies of heart failure.

## ACKNOWLEDGEMENTS AND FUNDING SOURCES

This work was supported by NIH grants R01-HL132111 and HL158671 (MT), R01-HL148598 (OK), R01-CA191785 (LX and JA), R01-CA243445 (LX), R33-CA252158 (LX), and American Heart Association Career Development Award CDA34110117 (OK). SS was supported by American Heart Association Predoctoral Fellowship (PRE35230020). LCG and was supported by an NIH Training grant T32HL125204 (PIs: Molkentin and Kranias) and American Heart Association Predoctoral Fellowship (PRE35210795).

