## Supplementary figures and images for "HuR inhibition reduces post-ischemic cardiac remodeling by dampening acute inflammatory gene expression and the innate immune response"

### Figure S1

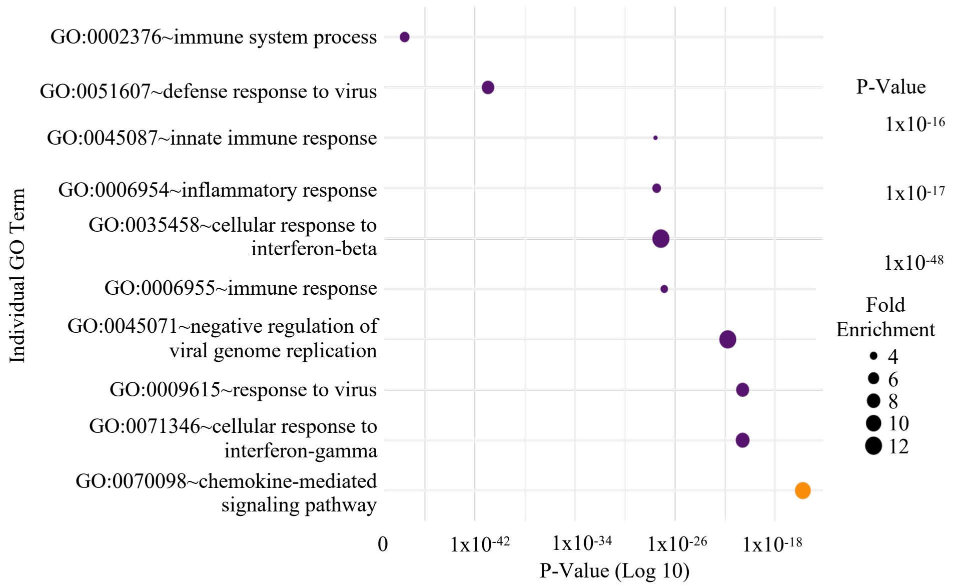

### Figure S2

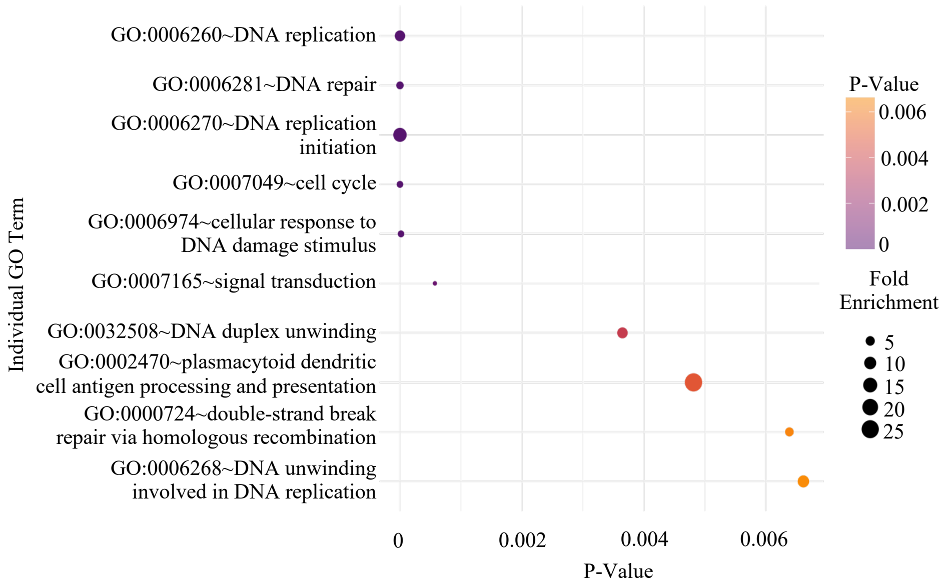

### Figure S3

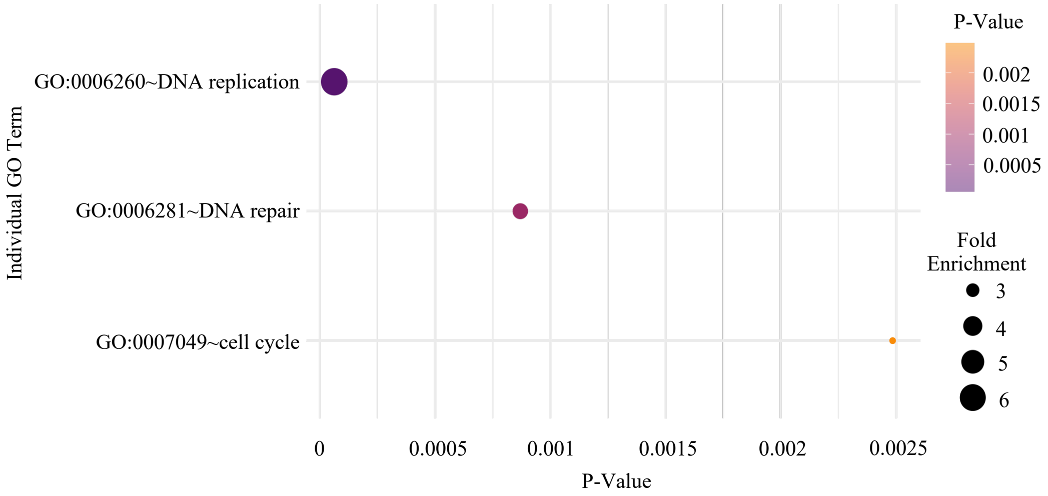
